# Diurnal rhythms with sunrise-induced oscular contractions suggest photosensitivity in the demosponge *Halichondria panicea*

**DOI:** 10.1101/2025.09.23.677974

**Authors:** Josephine Goldstein, Sarah B. Flensburg, Anders Garm, Peter Funch

## Abstract

Despite their quiescent lifestyle, sponges display contraction-expansion behaviours that regulate water flow through their aquiferous system. These behaviours are intrinsically generated but can be modulated by environmental parameters, including light exposure. Some sponges exhibit diurnal contraction-expansion rhythms synchronized with illumination cycles. Since the often-used photopigment opsin is unknown in sponges, the light-induced behaviour is likely mediated by cryptochromes and extra-ocular photoreceptors, as shown in sponge larvae. Previous studies on demosponges reported higher contraction-expansion activity during daylight than at night.

Understanding such diurnal behaviours in nerve- and muscle-less metazoans may provide insights into the early evolution of sleep-like states. However, systematic diurnal observations of sponge contraction-expansion rhythms remain scarce. Here, we document the contractile behaviour of single-osculum explants of the marine demosponge *Halichondria panicea* over six days, maintained under a 12:12 h dark:light cycle with simulated sunrise and sunset. We find a light-induced diurnal rhythm in the contraction-expansion behaviour of their osculum, suggesting photosensitivity.

Furthermore, we observe that sunrise triggers the onset of oscular contractions and discuss these results in context with the characteristics of sleep-like states in other metazoans.

**Summary statement:** A diurnal rhythm of light-induced oscular contractions suggests photosensitivity in the nerveless demosponge *Halichondria panicea.* This may offer insights into the early evolution of sleep-like behaviours in metazoans.

## Introduction

Sponges (poriferans) are filter-feeders which pump water through their aquiferous systems by means of flagellated choanocytes while retaining small particles such as free-living bacteria and phytoplankton cells with sizes down to 0.1 µm (Lüskow et al., 2019; Riisgård and Larsen, 2010). Water flow can be temporarily stopped by contractile behaviour (Goldstein et al., 2019; Kumala et al., 2017; Ludeman et al., 2017; Reiswig, 1971; Riisgård et al., 2016), which involves constriction of the inhalant openings (ostia), aquiferous canals, choanocyte chambers, as well as the exhalant openings (oscula) (Goldstein et al. 2020; Hammel and Nickel, 2014). Since sponges lack neurons and muscle cells, contractions seem to be mediated by a neuron-like toolkit of sensory cells, neurotransmitter-based signaling pathways and ionotropic receptor systems (Leys, 2015; Leys et al., 2019). Contractions can occur seemingly spontaneously in undisturbed sponges (Goldstein et al., 2020; Reiswig, 1971). They can also occur in response to environmental parameters such as resuspended sediment during storm events (Bell et al., 2015; Leys, 2015; Reiswig, 1971; Strehlow et al., 2017), mechanical disturbances (Leys, 2015; Leys and Mackie, 1997), oxygen depletion (Leys and Kahn, 2018) or the diurnal cycle of illumination (Reiswig, 1971).

Diurnal rhythms of contraction-expansion behaviour have so far been observed in two marine demosponges, *Tethya wilhelma* and *Tectitethya crypta* (Flensburg et al., 2022; Nickel, 2004; Reiswig, 1971). In both cases, the diurnal rhythm was seen as more active contraction-expansion behaviour during the day than during the night, indicating that sponges might show a sleep-like state during the quiescent period at night. This indication is highly interesting, since it could shed light on the early evolution of sleep (Keene and Duboue, 2018; Pennisi, 2021). Previous research has shown that sleep is present in animals with a well-developed brain such as chordates, arthropods, nematodes, molluscs, and platyhelminthes (Anafi et al., 2019; Pennisi, 2021). However, behaviourally defined sleep-like states (Lesku et al., 2009) have also been detected in cnidarians that are characterized by less centralized nervous systems, e.g. in the upside-down jellyfish *Cassiopea* based on its bell pulsing behaviour (Nath et al., 2017) and in the freshwater polyp *Hydra* based on tentacle movement (Kanaya et al., 2020). These cnidarian studies support the assumption that sleep arose early in the metazoan lineage, potentially before the nervous system.

Light seems to affect diurnal rhythms and behaviour in sponges, and both *Tethya wilhelma* and *Tectitethya crypta* increase their body contraction frequency under daylight conditions compared to nighttime (Reiswig 1971; Flensburg et al. 2022). Circadian clock genes may be involved in the regulation of this behaviour since in another demosponge, *Amphimedon queenslandica,* two clock genes exhibit an oscillating diurnal expression profile, and this rhythmic expression is partially lost when the animals are exposed to constant light or darkness (Jindrich et al., 2017). There are also indications of a more direct behavioural response to changes in the ambient light conditions, though, as *T. wilhelma* displays a much higher probability of contractions just after sunrise (Flensburg et al., 2022). The direct light control is also suggested by the diurnal rhythm disappearing straight away when the animals are kept in either constant light or constant darkness throughout the 24 hours of the day (Flensburg et al., 2022). Unlike other animals, sponges do not express the photosensitive opsin proteins and appear to utilize a cryptochrome-based photoreceptor system with non-retinal chromophores instead (Rivera et al., 2012).

Under the assumption that oscular contractions represent an active behaviour, and that periods with reduced contraction frequency represent a state of quiescence, we document the contraction-expansion behaviour of explants of the marine demosponge *Halichondria panicea* to gain further insight into the diurnal rhythms in sponges and their potential sleep-like behaviours.

## Materials and methods

### Cultivation of sponge explants

Sponges (*Halichondria panicea* Pallas, 1766) were collected from a stone wall at 0.5 m water depth in the inlet to Kerteminde Fjord, Denmark (55°26’59"N, 10°39’40"E) in July 2023 and brought to the nearby laboratory of the Marine Biological Research Centre. Small sponge pieces (∼0.5×0.5 cm, *n* = 100) were cut from the collected sponge material and placed on standard glass slides in a flow- through seawater tank (130 L, flow rate ∼1000 L d^‒1^) with sequentially filtered (200µm/80 µm) and UV-sterilized (EHEIM reflex UV 800) seawater, adjusted to stable water temperature (TECO TK 150 Tank Chiller Line) in a temperature-controlled room (15 °C). After 10 days, 83 sponge pieces had formed explants attached to the glass slides in the cultivation tank and regenerated functional aquiferous systems with a single osculum per explant (cf. Kumala et al. 2017). The light conditions in the cultivation tank were gradually adjusted from the ambient daylength of 17.5 h to a 12:12 h dark:light cycle by decreasing daylight by 30 minutes per day over 10 days. Illumination was provided by an LED lamp (FLUVAL Plant Spectrum), mounted about 30 cm above the sponge explants and controlled via the Fluval Smart App. The 12:12 h dark:light cycle consisted of night (light intensity <0.15 mW m^−2^) from 18:00 to 06:00, and day (light intensity 51 W m^−2^ at the water surface, see Fig. S1A for spectral properties) from 06:00 to 18:00. The light regime included a simulated sunrise from 06:00 to 07:00 and sunset from 17:00 to 18:00, during which light intensity increased and decreased linearly, respectively. Single-osculum explants were regularly fed heat-killed cells (jet function of EASYTRONIC MD112 microwave for 5 min) of the cryptophyte *Rhodomonas salina* Hill and Wetherbee, 1989 (NIVA-15/12, BlueBioTech GmbH, Germany; cultivated in the same temperature-controlled room under constant light conditions at a salinity of 20 psu using Walne’s medium, cf. Endar et al., 2012) by means of a peristaltic pump to resemble the chlorophyll *a* concentration of the sampling site in the inlet to Kerteminde fjord (where 799 *R. salina* cells mL^‒1^ correspond to 1 µg Chl *a* L^‒1^, cf. Riisgård et al., 2013).

### Video-observation of sponge explants

After an adaptation period of ≥21 days under defined light conditions, actively pumping sponge explants, as indicated by expanded oscula, were placed vertically in an open observation chamber in the cultivation tank where they were acclimated to the experimental conditions for 24 h. In addition to the LED lamp described above, an infrared (IR) lamp (HD Infrared Waterproof, see Fig. S1B for spectral properties) was placed at a similar height above the sponge explants in the observation chamber and provided continuous IR-light throughout the experiment (light intensity 140 W m^−2^ measured at the water surface). Side-view video-recordings of 16 randomly selected sponge explants in four experimental sets with four explants per set were obtained for a continuous period of six days using two digital camcorders (Panasonic HC-VXF1) operated in IR-sensitive night-shot mode. Water temperature (*T*, °C), salinity (*S*, psu) and chlorophyll *a* concentration (Chl *a*, µg L^‒1^) were monitored in the cultivation tank at the same diurnal interval with a YSI650 Multiparameter Display System. The same parameters were monitored daily at 12:00 pm and 24:00 pm at the sampling site in Kerteminde harbour (Table S1).

### Processing of video material

The videos were converted to time-lapse image sequences (interval: 1 min) using the software ffmpeg (version: 2023-02-27-git-891ed24f77-full_build, www.gyan.dev), and these images were used for all further analyses. Contractions of sponge explants were identified based on complete osculum closure and subsequent reduction in the explant projected area. Using imaging software (ImageJ, version 2.1.0), we measured the osculum diameter (*D*) to estimate the osculum cross-sectional area (*OSA* = π(*D*/2)², mm²; cf. Goldstein et al., 2019) along with the sponge projected area (*A*, mm²). We documented the timing and duration of contractile behaviour as defined by osculum closure, i.e. reduction of the initial (i.e., before contraction) *OSA* to zero during the contraction phase (I), followed by a contracted phase (II) with completely closed osculum and a subsequent expansion phase (III; cf. Goldstein et al., 2020). For comparison across treatments, we normalized all data setting maximum values to 1. Based on counts of the total number of osculum closure events (*N*) of each examined sponge explant (*n* = 16) during dark and light periods throughout the six day-observation period, we estimated the contraction frequency (*f*, [12 h]^‒1^) for each 12 h dark and light interval. We also calculated the proportion of oscular contractions (*p*, d^‒1^) for every hour within the 12:12 h dark:light cycle relative to the total number of osculum closure events throughout the six day-experimental period. We further calculated the time lag between osculum closure and reduction of the relative projected area to a minimum (Δ*t_OSA_*_‒*V*_, min) based on changes in osculum cross-sectional area (Δ*t_OSA_*, min) and projected area (Δ*t_A_*, min). Furthermore, we used these parameters to calculate the speed of the following processes: osculum closure (*v_OSA_*__I_, Δ*OSA*_r_ min^‒1^), osculum expansion (*v_OSA_*__III_, Δ*OSA*_r_ min^‒1^), and contraction and expansion of projected area (*v_A_*__I_/*v_A_*__III_, Δ*A_r_* min^‒1^).

### Statistical analyses

All statistical analyses were performed in R version 4.2.2 (R Core Team, 2020). Initially all response variables were checked for normality and homogeneity of variances by the Shapiro-Wilk test and F test, respectively. Differences in the total number of oscular contractions (*N*) and in the speed of osculum closure (*v_OSA_*__I_) or expansion (*v_OSA_*__III_;; Shapiro-Wilk normality test, p >0.05; F test, p >0.05; *n* = 16) in response to dark and light conditions were analyzed using one-way ANOVA. We further tested the timing of the first contraction after sunset or sunrise (Δ*t*_0_, min) and the duration of osculum closure (Δ*t_OSA_*__II_) for differences in response to dark and light conditions using generalized linear models (GLMs) with a Gamma error structure. The time lag between osculum closure and minimum projected area (Δ*t_OSA_*_‒*A*_), as well as contraction and expansion speeds of projected area (*v_A_*__I_ or *v_A_*__III_; Shapiro-Wilk normality test, p <0.05; F test, p <0.05; *n* = 16) in response to dark and light conditions were likewise tested with GLMs. Additional GLMs with Gamma error structure were parameterized to test for differences in maximum osculum cross-sectional area, water temperature (*T*), salinity (*S*) and chlorophyll *a* concentration (Chl *a*; Shapiro-Wilk normality test, p <0.05) between dark and light periods. This was also done to test for differences in the relative proportion of oscular contractions across sponge explants throughout the entire experimental period. Autocorrelation of residuals was examined using the Durbin-Watson test. We applied Pearson’s product-moment correlation to examine interdependence between the relative proportion of oscular contractions of individual sponge explants throughout the experiment. We further examined the relationship between contraction frequency and maximum osculum cross-section area (*OSA*_max_) by fitting a power function (i.e., a linear model, LM).

## Results

### Oscular contractions

All 16 explants displayed a dynamic osculum. When fully expanded, they had an osculum cross-sectional area (*OSA*_max_) of 1.01±0.68 mm^2^ (mean ± SD), and this size did not differ between dark and light periods (GLM, t = 0.64, p = 0.527). Throughout the experiment, we observed a total of 162 osculum closure events which followed a three-phase pattern (Figs. 1 and S2). Phase I, the oscular contraction phase, had a mean duration of 34±8 min. This was followed by phase II, a contracted state with the osculum remaining closed for 22±10 min. Finally, phase III, the oscular expansion phase (cf. Goldstein et al. 2020), lasted 19±9 min (Table 1 and Fig. 2). Oscular contractions were followed by slight reductions in sponge projected area to 98.2±0.9 % of the initial size (miminal size = 90.8 % of initial projected area; cf. Fig. 3). These minimum sizes of the explants were reached with a time lag of 11±8 min following oscular contractions (Table 2 and Fig. 3). The oscula of sponge explants contracted 3.4±2.3 times during dark periods (0 to 7 contractions) and doubled to 6.8±1.6 (3 to 11) contractions during light periods throughout the six day-experiment, resulting in contraction frequencies of 0.6±0.4 per 12 h-dark interval and 1.1±0.3 per 12 h-light interval, respectively (Table 1). There were significantly more oscular contractions during light compared to dark periods (ANOVA, F_1,30_ = 20.91, p <8×10^‒5^; Fig. 4A). The timing of the first osculum closure events after sunrise at 06:00 am, 2.2±1.1 h, was significantly shorter than after sunset at 18:00 pm, 6.2±1.9 h (GLM, t = ‒5.64, p <5×10^‒5^, Table 1, Fig. 4B). There was a strong correlation between sunrise and the initiation of oscular contractions, with a high proportion of contractions (41 %) happening between 06:00 am and 09:00 am, i.e., within 3 h after sunrise (Fig. 5). Relative proportions of oscular contractions showed no correlation between sponge explants (Pearson’s product-moment correlation, t_2110_ <4×10^‒17^, p = 0.500), remained constant throughout the six-day experimental period (GLM, t = ‒0.31, p = 0.755), and exhibited no autocorrelation in the residuals (Durbin Watson test, DW = 2.11, p = 0.996).

**Figure 1.**
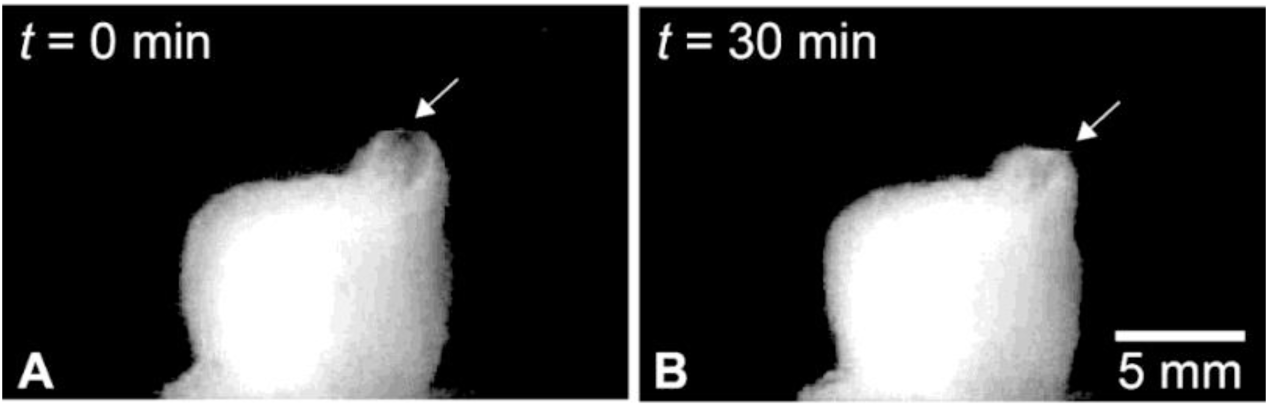
Contraction of the osculum in *Halichondria panicea*. Sponge explant (ID #15) A) with an expanded osculum at time *t* = 0 min and B) with a contracted osculum at *t* = 30 min. Arrows indicate the position of the osculum in both states.

**Figure 2.**
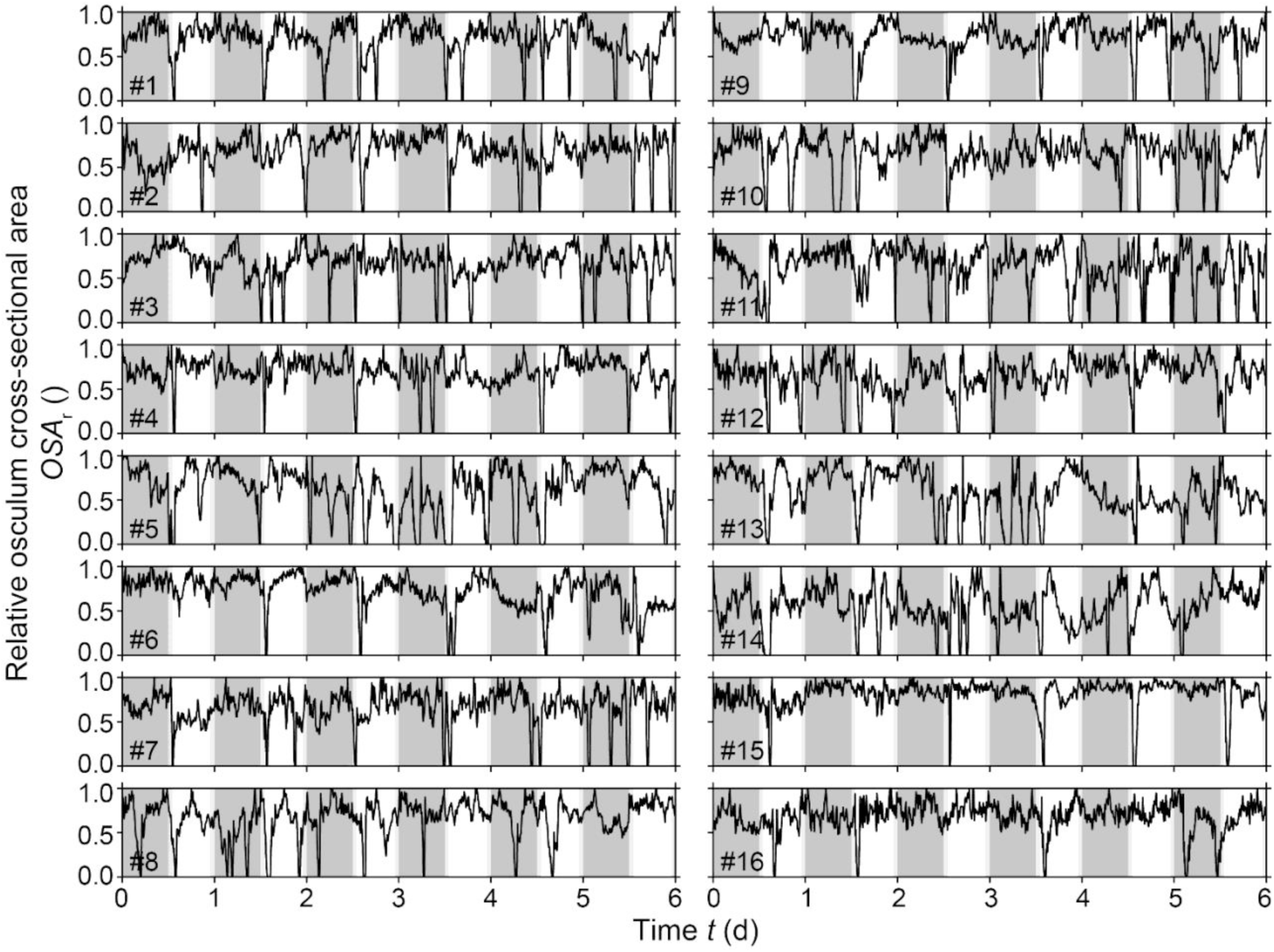
Dynamics of the osculum cross-sectional area (*OSA*_r_) of sponge explants (*n* = 16; ID #1‒ 16) over a six-day observation period under controlled 12-hour dark/light cycles. The diagram shows dark periods (dark grey), light periods (white), and transitions at sunrise and sunset (light grey). Note the distinct relatively short-term events where the oscula close completely, i.e., reduction of *OSA*_r_ to zero, and subsequently expand.

**Figure 3.**
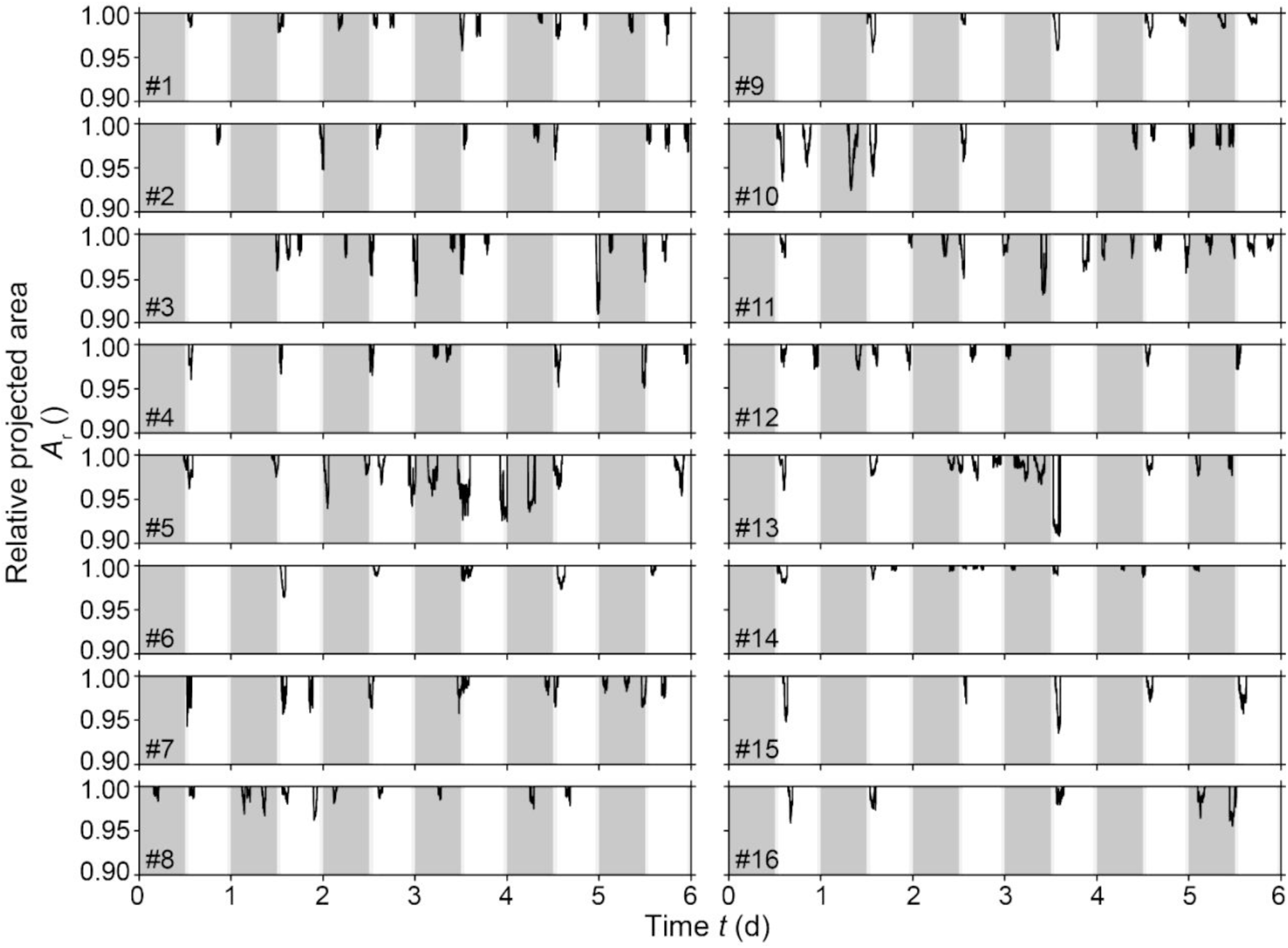
*Halichondria panicea*. Relative projected area (*A*_r_) of sponge explants (*n* = 16; ID #1‒16) throughout the six-day observation period under controlled light conditions following a 12:12 h dark:light cycle. Dark periods (dark grey areas) and light periods (white areas), including sunrise and sunset (light grey areas), are shown. Osculum closure is followed by subsequent changes in projected area, i.e., reductions by <10 %, and subsequent expansion of the sponge explant.

**Figure 4.**
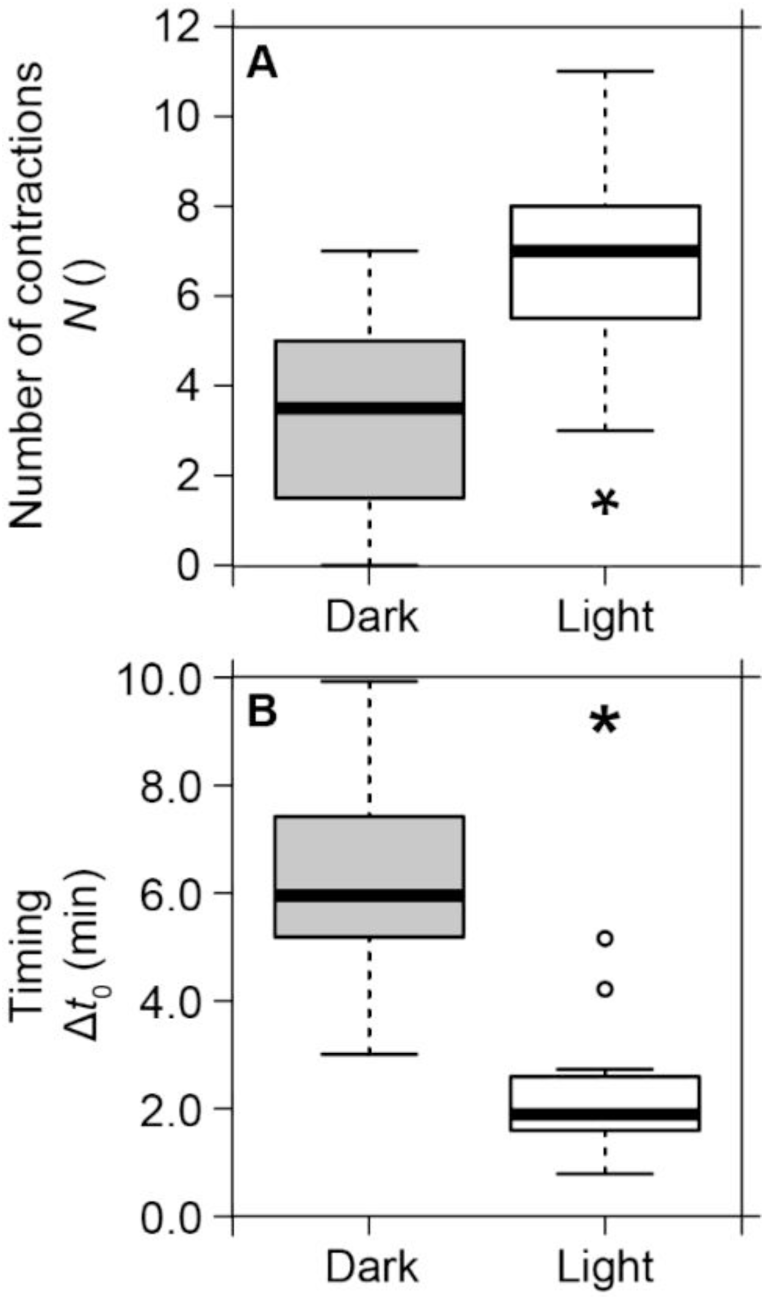
*Halichondria panicea*. Box-and-whisker plots of A) total oscular contractions throughout the experimental period and B) time to first oscular contraction after sunset and sunrise, respectively, during dark (grey) versus light (white) periods. Based on the total number of complete osculum closure events observed in sponge explants (*n* = 16) kept at a 12:12 h dark:light cycle for six days. Medians (solid black lines), lower (25 %) and upper (75 %) quartiles (boxes) are shown along with minimum and maximum values (whiskers), as well as outliers (open circles). Asterisks indicate significant differences between dark and light periods (p <0.05). Based on data from Table 1.

**Figure 5.**
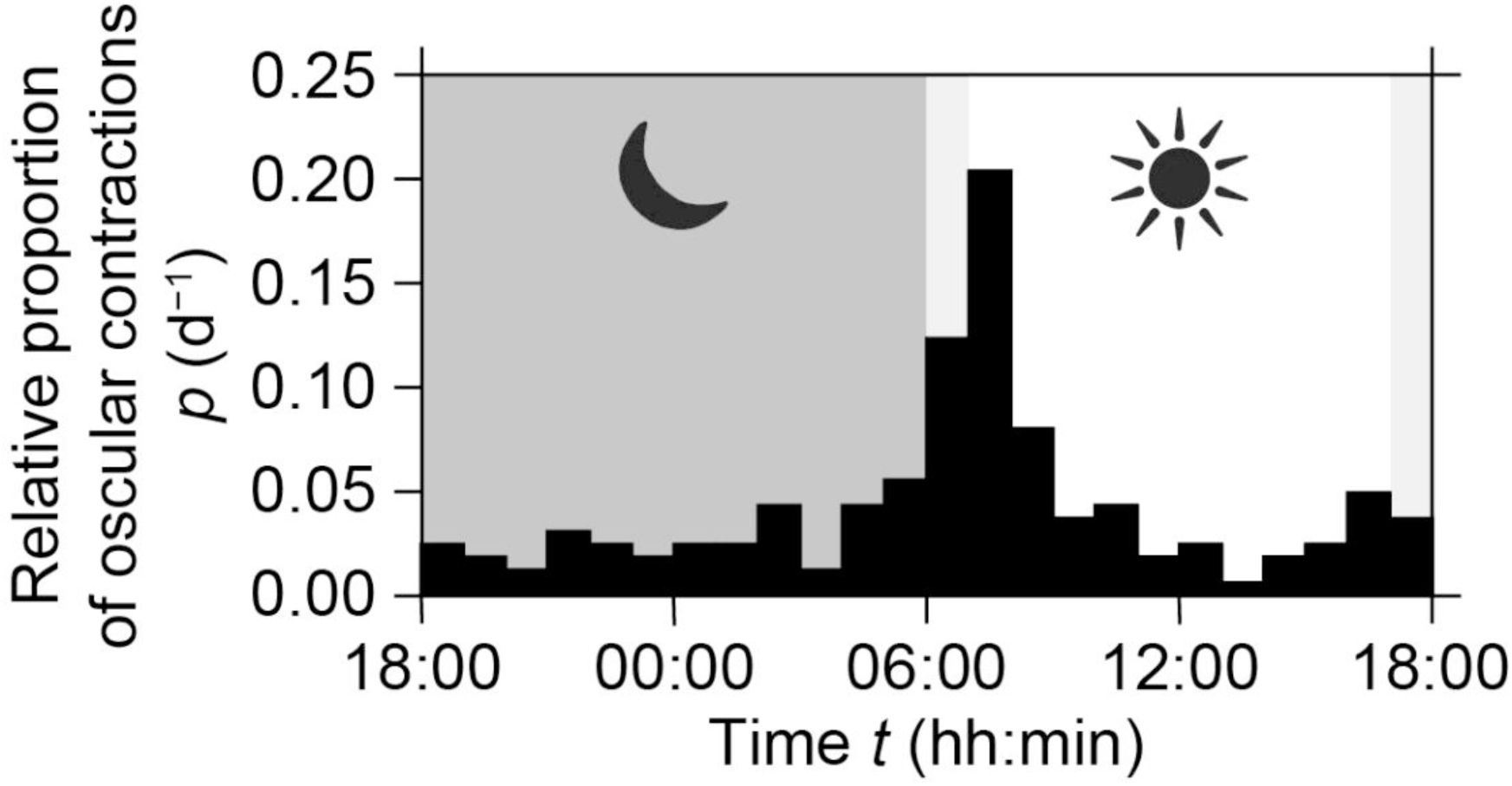
Mean relative proportions of contractions of sponge explants (*n* = 16) with a 1 h time resolution across the 12:12 h dark:light cycle, based on a total of 162 contractions throughout the six-day observation period. Dark grey indicates dark periods (nighttime), white indicates light periods (daytime), and light grey indicates the transitions at sunrise and sunset.

**Table 1.** Contractile behaviour of *Halichondria panicea* (ID #1‒16) kept at a 12:12 h dark:light cycle (grey:white) for six days. ID #: sponge explant number; *N*: total number of contractions during dark and light intervals throughout the experimental period; *f* [= *N*/6)]: contraction frequency per 12 h dark:light interval; Δ*t_0_*: timing of first contraction after sunset or sunrise, respectively; Δ*t_OSA_*: duration of phase of oscular contraction (I), contracted phase (II) and phase of oscular expansion (III); *OSA*: initial osculum cross-sectional area (I, II and III); *OSA*_r_ (= *OSA*/*OSA*_max_): relative osculum cross-sectional area (I, II and III); *v_OSA_* (= Δ*OSA*_r_ /Δ*t_OSA_*): speed of osculum closure (I) and expansion (III). Means (±SD) are shown.

| Phase | ID # | $N$ () | $f$ (12 h)<sup>-1</sup> | $\Delta t_0$ (min) | $\Delta t_{OSA}$ (min) | | | $OSA$ (mm<sup>2</sup>) | | | $OSA_r$ () | | | $v_{OSA}$ ($\Delta OSA_r \text{ min}^{-1}$) | |
| --- | --- | --- | --- | --- | --- | --- | --- | --- | --- | --- | --- | --- | --- | --- | --- |
|  |  |  |  |  | I | II | III | I | II | III | I | II | III | I | III |
| Dark | 1 | 3 | 0.5 | 7.2±2.2 | 30±10 | 15±10 | 12±6 | 0.38 | 0 | 0.42 | 0.90 | 0 | 1.00 | -0.030 | 0.083 |
| | 2 | 1 | 0.2 | 7.4±0.0 | 25±0 | 39±0 | 14±0 | 0.45 | 0 | 0.81 | 0.56 | 0 | 1.00 | -0.022 | 0.071 |
| | 3 | 6 | 1.0 | 9.9±12.3 | 27±11 | 14±2 | 11±3 | 0.55 | 0 | 0.62 | 0.89 | 0 | 1.00 | -0.033 | 0.091 |
| | 4 | 2 | 0.3 | 5.6±0.0 | 40±4 | 17±2 | 13±7 | 0.72 | 0 | 0.85 | 0.85 | 0 | 1.00 | -0.021 | 0.077 |
| | 5 | 5 | 0.8 | 5.8±4.5 | 35±16 | 34±25 | 17±2 | 0.67 | 0 | 0.72 | 0.93 | 0 | 1.00 | -0.027 | 0.059 |
|  | 6 | 0 | 0.0 | - | - | - | - | - | - | - | - | - | - | - | - |
| | 7 | 5 | 0.8 | 7.9±5.6 | 26±3 | 21±5 | 17±4 | 0.73 | 0 | 0.73 | 1.00 | 0 | 1.00 | -0.038 | 0.059 |
| | 8 | 7 | 1.2 | 4.8±1.6 | 28±10 | 14±4 | 14±3 | 0.50 | 0 | 0.80 | 0.63 | 0 | 1.00 | -0.023 | 0.071 |
| | 9 | 1 | 0.2 | 8.4±0.0 | 44±0 | 27±0 | 32±0 | 1.72 | 0 | 1.60 | 1.00 | 0 | 0.93 | -0.023 | 0.029 |
| | 10 | 5 | 0.8 | 6.1±4.7 | 33±4 | 29±38 | 21±7 | 0.82 | 0 | 0.92 | 0.90 | 0 | 1.00 | -0.027 | 0.048 |
| | 11 | 6 | 1.0 | 6.5±3.7 | 38±17 | 15±8 | 15±4 | 0.66 | 0 | 0.62 | 1.00 | 0 | 0.94 | -0.026 | 0.063 |
| | 12 | 2 | 0.3 | 5.4±6.5 | 41±16 | 17±1 | 15±7 | 0.64 | 0 | 0.79 | 0.81 | 0 | 1.00 | -0.020 | 0.067 |
| | 13 | 5 | 0.8 | 3.0±1.1 | 49±22 | 43±41 | 21±4 | 0.59 | 0 | 0.49 | 1.00 | 0 | 0.82 | -0.020 | 0.039 |
| | 14 | 4 | 0.7 | 5.2±4.0 | 25±6 | 20±6 | 15±5 | 0.41 | 0 | 0.59 | 0.70 | 0 | 1.00 | -0.028 | 0.067 |
|  | 15 | 0 | 0.0 | - | - | - | - | - | - | - | - | - | - | - | - |
| | 16 | 2 | 0.3 | 3.1±0.0 | 38±6 | 12±8 | 55±7 | 2.75 | 0 | 1.89 | 1.00 | 0 | 0.69 | -0.026 | 0.013 |
| Light | 1 | 9 | 1.5 | 1.8±1.9 | 27±6 | 16±6 | 16±7 | 0.41 | 0 | 0.45 | 0.92 | 0 | 1.00 | -0.034 | 0.063 |
| | 2 | 8 | 1.3 | 4.2±4.7 | 23±4 | 18±4 | 13±4 | 0.57 | 0 | 0.68 | 0.84 | 0 | 1.00 | -0.037 | 0.077 |
| | 3 | 7 | 1.2 | 1.5±2.3 | 25±4 | 19±6 | 11±3 | 0.56 | 0 | 0.51 | 1.00 | 0 | 0.92 | -0.040 | 0.084 |
| | 4 | 6 | 1.0 | 0.8±0.6 | 28±5 | 16±10 | 14±2 | 0.70 | 0 | 0.70 | 1.00 | 0 | 0.99 | -0.036 | 0.071 |
| | 5 | 8 | 1.3 | 2.7±3.9 | 39±24 | 49±31 | 19±6 | 0.53 | 0 | 0.57 | 0.94 | 0 | 1.00 | -0.024 | 0.053 |
| | 6 | 6 | 1.0 | 1.8±0.6 | 41±16 | 15±6 | 21±8 | 1.57 | 0 | 1.30 | 1.00 | 0 | 0.83 | -0.024 | 0.040 |
| | 7 | 7 | 1.2 | 1.7±1.5 | 31±9 | 13±2 | 19±9 | 0.73 | 0 | 0.68 | 1.00 | 0 | 0.93 | -0.032 | 0.049 |
| | 8 | 5 | 0.8 | 2.6±1.0 | 28±8 | 25±15 | 16±6 | 0.52 | 0 | 0.71 | 0.72 | 0 | 1.00 | -0.026 | 0.063 |
| | 9 | 6 | 1.0 | 2.1±1.9 | 52±26 | 22±13 | 27±12 | 1.74 | 0 | 1.61 | 1.00 | 0 | 0.92 | -0.019 | 0.034 |
| | 10 | 5 | 0.8 | 1.7±0.7 | 34±10 | 24±11 | 23±10 | 0.94 | 0 | 0.78 | 1.00 | 0 | 0.82 | -0.029 | 0.036 |
| | 11 | 11 | 1.8 | 5.2±4.2 | 31±17 | 20±10 | 16±8 | 0.65 | 0 | 0.68 | 0.96 | 0 | 1.00 | -0.031 | 0.063 |
| | 12 | 7 | 1.2 | 2.1±1.1 | 37±8 | 18±4 | 16±2 | 0.54 | 0 | 0.77 | 0.70 | 0 | 1.00 | -0.019 | 0.063 |
| | 13 | 7 | 1.2 | 1.5±0.8 | 45±13 | 29±15 | 21±9 | 0.78 | 0 | 0.87 | 0.89 | 0 | 1.00 | -0.020 | 0.048 |
| | 14 | 8 | 1.3 | 1.1±0.5 | 26±12 | 26±22 | 17±3 | 0.44 | 0 | 0.73 | 0.60 | 0 | 1.00 | -0.023 | 0.059 |
| | 15 | 5 | 0.8 | 2.0±0.5 | 36±14 | 12±5 | 27±12 | 2.21 | 0 | 2.46 | 0.90 | 0 | 1.00 | -0.025 | 0.037 |
| | 16 | 3 | 0.5 | 2.6±1.2 | 39±11 | 9±3 | 32±5 | 3.03 | 0 | 2.46 | 1.00 | 0 | 0.81 | -0.026 | 0.025 |

**Table 2.** Contractile behaviour of *Halichondria panicea* (ID #1‒16) kept at a 12:12 h dark:light cycle (grey:white) for six days. ID #: sponge explant number; *V*_m_: measured sponge volume; Δ*t_OSA‒ A_*: time lag between osculum closure and reduction in sponge projected area; Δ*t_A_*: duration of contraction (I) and expansion (III) of projected area; *A*_r_ (= *A*/*A*_max_): relative projected area with a maximum (*A*_max_) in phase I or III, and a minimum during the contracted phase (II); *v_A_* (= Δ*A*_r_ /Δ*t_A_*): contraction and expansion speeds (I and III) of projected area. Means (±SD) are shown.

| Phase | ID # | $V_m$<br>(mm<sup>3</sup>) | $\Delta t_{OSA-A}$<br>(min) | $\Delta t_A$<br>(min) | | $A_r$<br>( ) | | | $v_A$<br>($\Delta A_r \times 10^{-3} \text{ min}^{-1}$) | |
| --- | --- | --- | --- | --- | --- | --- | --- | --- | --- | --- |
|  |  |  |  | I | III | I | II | III | I | III |
| Dark | 1 | 177 | 4±7 | 35±9 | 23±9 | 1.000 | 0.991 | 0.999 | −0.257 | 0.348 |
| | 2 | 191 | 30±0 | 55±0 | 23±0 | 1.000 | 0.984 | 0.991 | −0.291 | 0.304 |
| | 3 | 199 | 4±4 | 31±13 | 21±5 | 1.000 | 0.966 | 0.990 | −1.097 | 1.143 |
| | 4 | 210 | 10±2 | 49±6 | 20±7 | 0.996 | 0.993 | 1.000 | −0.061 | 0.350 |
| | 5 | 276 | 6±4 | 41±16 | 45±26 | 1.000 | 0.976 | 0.989 | −0.585 | 0.289 |
|  | 6 | 277 | - | - | - | - | - | - | - | - |
| | 7 | 300 | 8±6 | 34±6 | 29±10 | 0.991 | 0.983 | 1.000 | −0.235 | 0.586 |
| | 8 | 334 | 9±4 | 37±11 | 19±6 | 1.000 | 0.988 | 0.997 | −0.324 | 0.474 |
| | 9 | 341 | 27±0 | 71±0 | 32±0 | 1.000 | 0.990 | 0.991 | −0.141 | 0.031 |
| | 10 | 348 | 9±13 | 42±12 | 40±32 | 1.000 | 0.976 | 0.992 | −0.571 | 0.400 |
| | 11 | 348 | 8±5 | 45±17 | 22±4 | 1.000 | 0.981 | 0.998 | −0.422 | 0.773 |
| | 12 | 363 | 3±1 | 44±15 | 29±5 | 0.998 | 0.985 | 1.000 | −0.295 | 0.517 |
| | 13 | 372 | 32±40 | 80±58 | 33±9 | 1.000 | 0.984 | 0.993 | −0.200 | 0.273 |
| | 14 | 470 | 9±6 | 34±10 | 26±10 | 1.000 | 0.997 | 0.999 | −0.088 | 0.077 |
|  | 15 | 524 | - | - | - | - | - | - | - | - |
| | 16 | 542 | 3±4 | 40±3 | 64±4 | 0.982 | 0.967 | 1.000 | −0.375 | 0.516 |
| Light | 1 | 177 | 9±6 | 36±9 | 23±12 | 1.000 | 0.989 | 1.000 | −0.306 | 0.478 |
| | 2 | 191 | 8±5 | 30±8 | 23±7 | 0.999 | 0.987 | 1.000 | −0.400 | 0.565 |
| | 3 | 199 | 7±5 | 32±6 | 23±3 | 1.000 | 0.978 | 0.998 | −0.688 | 0.870 |
| | 4 | 210 | 9±9 | 37±11 | 21±4 | 0.990 | 0.969 | 1.000 | −0.568 | 1.476 |
| | 5 | 276 | 18±13 | 57±27 | 50±26 | 1.000 | 0.968 | 0.993 | −0.561 | 0.500 |
| | 6 | 277 | 7±8 | 48±14 | 29±6 | 1.000 | 0.989 | 0.997 | −0.229 | 0.276 |
| | 7 | 300 | 7±5 | 38±9 | 24±7 | 0.997 | 0.985 | 1.000 | −0.316 | 0.625 |
| | 8 | 334 | 16±17 | 44±17 | 24±8 | 0.994 | 0.987 | 1.000 | −0.159 | 0.542 |
| | 9 | 341 | 17±15 | 69±28 | 33±7 | 1.000 | 0.984 | 0.991 | −0.232 | 0.212 |
| | 10 | 348 | 15±13 | 49±19 | 31±11 | 1.000 | 0.964 | 0.985 | −0.735 | 0.677 |
| | 11 | 348 | 11±11 | 42±21 | 25±15 | 1.000 | 0.983 | 0.998 | −0.405 | 0.600 |
| | 12 | 363 | 6±5 | 43±10 | 28±6 | 1.000 | 0.985 | 0.997 | −0.349 | 0.429 |
| | 13 | 372 | 17±18 | 62±12 | 32±15 | 1.000 | 0.980 | 0.989 | −0.323 | 0.281 |
| | 14 | 470 | 18±22 | 44±33 | 26±5 | 1.000 | 0.994 | 0.999 | −0.136 | 0.192 |
| | 15 | 524 | 8±6 | 45±16 | 31±11 | 1.000 | 0.966 | 0.978 | −0.756 | 0.387 |
| | 16 | 542 | 3±4 | 42±7 | 38±10 | 1.000 | 0.981 | 0.997 | −0.452 | 0.421 |

### Speed and duration of oscular closure and expansion

The speed of oscular closure ranged from ‒1.9 to ‒4.0 % Δ*OSA*_r_ min^‒1^ and was comparable during dark (‒2.6±0.5 % Δ*OSA*_r_ min^‒1^) and light periods (‒2.8±0.7 % Δ*OSA*_r_ min^‒1^; ANOVA, F_1,28_ = 0.77, p = 0.389). The duration of oscular closure in dark periods was 23±10 min, which was similar in light periods: 21±9 min (GLM, t = ‒0.54, p = 0.596). The speed of oscular expansion ranged from 1.2 to 8.8 % Δ*OSA*_r_ min^‒1^ and there was no significant difference between nighttime and daytime behaviour (dark: 5.4±1.7 % Δ*OSA_r_* min^‒1^, light: 6.0±2.1 % Δ*OSA*_r_ min^‒1^; ANOVA, F_1,28_ = 0.67, p = 0.421; Table 1). There was no significant diurnal pattern in the contraction speed of sponge projected area (dark: ‒0.035±0.026 % Δ*A*_r_ min^‒1^, light: ‒0.041±0.019 % Δ*A*_r_ min^‒1^; GLM, t = 390.7, p = 0.519). Also, we did not find a significant diurnal pattern in the time lag between complete osculum closure and minimum projected area (dark: 12±10 min, light: 11±5 min; GLM, t = ‒0.15, p = 0.880). Finally, the speed of expansion of projected area did not display a significant diurnal pattern either (dark: 0.043±0.027 % Δ*A*_r_ min^‒1^, light: 0.053±0.030 % Δ*A*_r_ min^‒1^; GLM, t = 403.8, p = 0.402; Table 2). The average osculum closure speed of ‒2.7±0.6 % Δ*OSA*_r_ min^‒1^ was significantly slower than oscular expansion speed of 5.7±1.9 % Δ*OSA*_r_ min^‒1^ (GLM, t =‒9.07, p <2×10^‒12^, Table 1), while contraction and expansion speeds of sponge projected area showed the similar mean values of ‒0.039±0.023 ΔAr h^‒1^ and 0.049±0.030 ΔAr h^‒1^, respectively (GLM, t = ‒528.0, p = 0.152, Table 2).

Water temperature (GLM, t = ‒0.14, p = 0.892), salinity (GLM, t = 0.34, p = 0.735) and Chl *a* concentration (GLM, t = ‒0.05, p = 0.962) in the cultivation tank remained stable throughout dark (*T* = 15.8±1.2 °C, *S* = 19.5±2.0 psu, Chl *a* = 7.9±1.6 µg L^‒1^) and light periods (*T* = 15.7±1.3 °C, *S* = 19.7±2.0 psu, Chl *a* = 7.8±1.6 µg L^‒1^). Furthermore, these conditions were comparable with the hydrography at the sampling site in Kerteminde harbour (*T* = 17.2±0.8 °C, *S* = 20.4±1.8 psu, Chl *a* = 8.1±1.3 µg L^‒1^; Table S1).

## Discussion

### Diurnal rhythms in contraction-expansion behaviour

The present work demonstrates a diurnal rhythm with reduced frequencies of osculum closure and subsequent small whole-body contractions during nights compared to daytime in the demosponge *Halichondria panicea*. Osculum closure in *H. panicea* is correlated with the onset of light, as seen by a clear peak in closures in the period 1-2 h after sunrise. Similar diurnal rhythms have previously been observed in *Tethya wilhelma* (Flensburg et al., 2022; Nickel, 2004) and in *Tectitethya crypta* (Reiswig, 1971). Interestingly, whole-body contraction-expansions also peaked just after sunrise in *Tethya wilhelma* (Flensburg et al., 2022), while reductions in pumping activity due to osculum closure followed sunset in *Tectitethya crypta* (Reiswig, 1971). Interpretation of this correlation with either sunrise or sunset is currently hampered by the fact that the function and purpose of sponge contraction-expansion behaviour are not finally settled. It has been suggested that the freshwater sponge *Ephydatia muelleri* performs peristaltic-like contractions that expel waste from its aquiferous system (Elliott and Leys, 2007), while in *H. panicea* high particle concentrations trigger osculum closure and body contractions, possibly with a similar purpose (Goldstein et al., 2024). In *Aplysina archeri*, mucus traps and transports particles against incoming flow, and contractions may function like a sneeze to eject debris (Kornder et al., 2022). Beyond preventing clogging, contraction-expansion behaviour may also play a role in supporting and controlling the activity of their internal microbiome (Kumala et al., 2021). This idea is reinforced by the finding that phototrophic symbionts can drive diurnal patterns in the sponge holobiont either through their own rhythmic activity or by serving as a food source (Sacristán-Soriano et al., 2019). For example, energy acquisition in the demosponge *Cliona orientalis* depends on a marked day-night relocation of its symbiotic dinoflagellates (Fang et al., 2016). Species in the genus *Halichondria* are generally low microbial abundance (LMA) sponges, exhibiting high variability in bacterial diversity across species and environments (reviewed by Goldstein and Funch, 2022; Knobloch et al., 2019; Sacristán-Soriano et al., 2019). The potential role of symbiotic interactions with phototrophic organisms, such as cyanobacteria occasionally found in *H. panicea*, remains unclear. However, the core microbiome of *H. panicea* is dominated by the heterotrophic alphaproteobacterium ‘*Candidatus* Halichondribacter symbioticus’ (Imhoff and Stöhr, 2003; Knobloch et al., 2019, 2020; Vethaak et al., 1982), and it is not yet known whether this bacterium exhibits any diurnal activity patterns.

The rhythmicity of sponge contraction-expansion behaviour has been shown to be strongly influenced by external factors. Periods of prolonged osculum closure in response to sediment disturbance have previously been described as a longer-term ‘dormant’ state in *Ianthella basta* (Strehlow et al., 2017), while the circadian behavioural cycle of *T. crypta* populations reveals seasonal differences with reduced pumping activity during storms in winter compared to calm summer periods (Reiswig, 1971, 1974). Thus, diurnal contraction rhythms of sponges can be overruled by environmental disturbances, including other factors such as seasonal temperature changes, biotic interferences through predation or mechanical disturbance, e.g. by calanoid copepods (Reiswig, 1974). Diurnal rhythms can also be influenced by release of gas bubbles by ingested phototrophic microorganisms, or accumulation of waste material on the sponge surface (Goldstein et al., 2024), as observed in a preliminary setting of the present work.

### Light influence on sponge contraction-expansion behaviour

In the present study, we show that the oscular contractions of *H. panicea* are induced by light, while the underlying mechanisms are still unknown. Previous findings for *T. wilhelma* show that the diurnal contraction rhythms for this sponge species are driven directly by the ambient light intensity, since rhythmicity disappears when it is kept under constant light or dark conditions (Flensburg et al., 2022). In *Amphimedon queenslandica,* rhythmic diurnal expression of the putative circadian clock genes is disrupted under constant light or darkness (Jindrich et al., 2017). Both systems strongly indicate that photosensitivity is involved in regulating the diurnal rhythms of sponges. However, the typical metazoan photopigment, opsin, appears to be absent in sponges (Feuda et al. 2012). Instead, evidence suggests that alternative photopigments such as cryptochromes or cytochromes mediate their light-guided behaviours (Jindrich et al., 2017; Lin and Todo, 2005; Rivera et al., 2012). Support for this includes the action spectrum of *Haliclona* sp., which closely matches the absorption spectrum of reduced cytochrome *c* oxidase (Björn and Rasmusson, 2009), and findings that sponge larvae exhibit peak sensitivity to blue light (440 nm) with a secondary peak in orange-red light (600 nm), suggesting a flavin- or carotenoid-based photopigment (Leys et al., 2002). Furthermore, cryptochrome expression in *Suberites domuncula* (tissue and primmorphs) is increased after exposure to light (Wang et al., 2012) and a blue-light-receptive cryptochrome with a flavin-based co-factor has been identified in the pigment ring of *A. queenslandica* larvae (Rivera et al., 2012).

### Frequency of sponge contraction-expansion behaviour

The observed contraction and expansion frequencies for osculum closure and subsequent changes in sponge projected area are in good agreement with previous findings for undisturbed *H. panicea* explants and their neurotransmitter- or particle-induced contractions (Goldstein et al., 2020; their Tables 2 and 3). Our data demonstrates a significant inverse relationship between the contraction frequency of *H. panicea* and the maximum (expanded) osculum cross-sectional area (*OSA*_max_), following the power function *f* = 1.51*OSA*_max_^‒0.60^ (LM, t = ‒7.20, p <5×10^‒6^; Fig. S3). This finding implies that observed contraction frequencies of 0.8 to 2.8 d^-1^ may be even lower under natural conditions, where sponges grow multiple larger oscula, each corresponding to an aquiferous module of a certain maximum size and maximum filtration rate (Goldstein and Funch, 2022; Kealy et al., 2019; Riisgård et al., 2024). In addition, larger *Halichondria* may possess static, permanently open pseudo-oscula (cf. Riisgård and Larsen 2022), challenging field observations on the diurnal rhythms of sponge species of similar morphology. Much higher contraction frequencies of 4 to 8 d^-1^ have been observed in *T. wilhelma* of similar size (0.5 to 1.0 cm) as the *H. panicea* explants used here (Flensburg et al., 2022). This pronounced difference could be a mere species difference but could also be influenced by temperature since *T. wilhelma* is thought to originate from tropical waters (even though only known from aquaria) and was tested at 22 °C (Flensburg et al., 2022), whereas the temperate *H. panicea* were tested at ∼15 °C in the present study.

### Do quiescent sponges sleep?

The present findings could indicate a quiescent state with reduced activity and lower energy consumption at night for *H. panicea*. Fewer contractions at night potentially result in more time actively pumping, but the energetic cost for pumping is generally low in demosponges (<1 % of the entire metabolism, cf. Riisgård et al., 1993; but see Hadas et al. 2008, Leys et al. 2011, Ludeman et al. 2017) and continuous beating of choanocyte flagella has been observed after inducing contractions in *H. panicea* (Riisgård et al., 2023, cf. Funch et al., 2023). New data from the freshwater sponge *Spongilla lacustris* might turn the picture around, though. Here, it was found that contrary to previous studies that interpreted body volume reductions in the marine sponge *T. wilhelma* as contractions of the epithelial cells (Nickel et al., 2011), contractions may actually be due to the relaxation of actomyosin stress fibers in epithelial canal cells (Ruperti et al., 2024). This would mean that *H. panicea* is in a quiescent state during daytime, but any conclusions await a better understanding of the energy demands of the sponge contraction-expansion behaviour.

However, independent of whether it is at nighttime or during the day that sponges like *H. panicea* and *T. wilhelma* are quiescent, the observed diurnal behaviour has the exciting possibility of being an evolutionary precursor of sleep.

Recent work has characterized the diurnal behaviours of the upside-down jellyfish *Cassiopea* sp. and the freshwater polyp *Hydra vulgaris* as sleep based on these behaviours being reversible quiescence at night, with reduced responsiveness to stimuli during the quiescent state and with homeostatic regulation (Nath et al., 2017; Kanaya et al., 2020). With these results, there is evidence that all animals with a nervous system display sleep-like behaviours, begging the question whether sleep evolved as a consequence of nervous systems or whether it evolved even earlier (Nath et al., 2017). We propose that sponges are the right animals to answer this question, and with our results documenting their diurnal rhythms, including a quiescent state, we have laid the ground for testing if this represents a sleep-like state. Finding the right behavioural paradigm to test for sleep-like responses is now the major challenge, not least since all reactions in sponges happen extremely slowly. Still, we believe that the contraction-expansion behaviour is promising, especially as it is possible to provoke this behaviour with physical disturbance (Reiswig, 1974). The hypothesis is that *Halichondria panicea* will respond less to physical disturbances when quiescent and that they will prolong their quiescent state with repeated disturbances. If this hypothesis is true, then it would suggest that the nerve- and muscle-less sponges sleep and lead to the surprising conclusion that sleep outdates the nervous system in animals.

## Acknowledgements

We are grateful to Fjord & Bælt for kindly providing access to the sampling site in Kerteminde harbour, and Majken Kristina Gjørup-Olesen for fruitful discussions on the diurnal behaviour of sponges.

## Author contributions

Conceptualization: J.G., S.B.F., A.G., P.F.; Methodology: J.G., S.B.F, A.G., P.F.; Formal analysis: J.G.; Investigation: J.G.; Resources: P.F.; Writing - original draft: J.G.; Writing - review and editing: J.G., S.B.F., A.G., P.F.; Visualization: J.G.; Funding acquisition: A.G., P.F.

## Data availability

All relevant data can be found within the article and its supplementary information.

---

**Figure S1.**
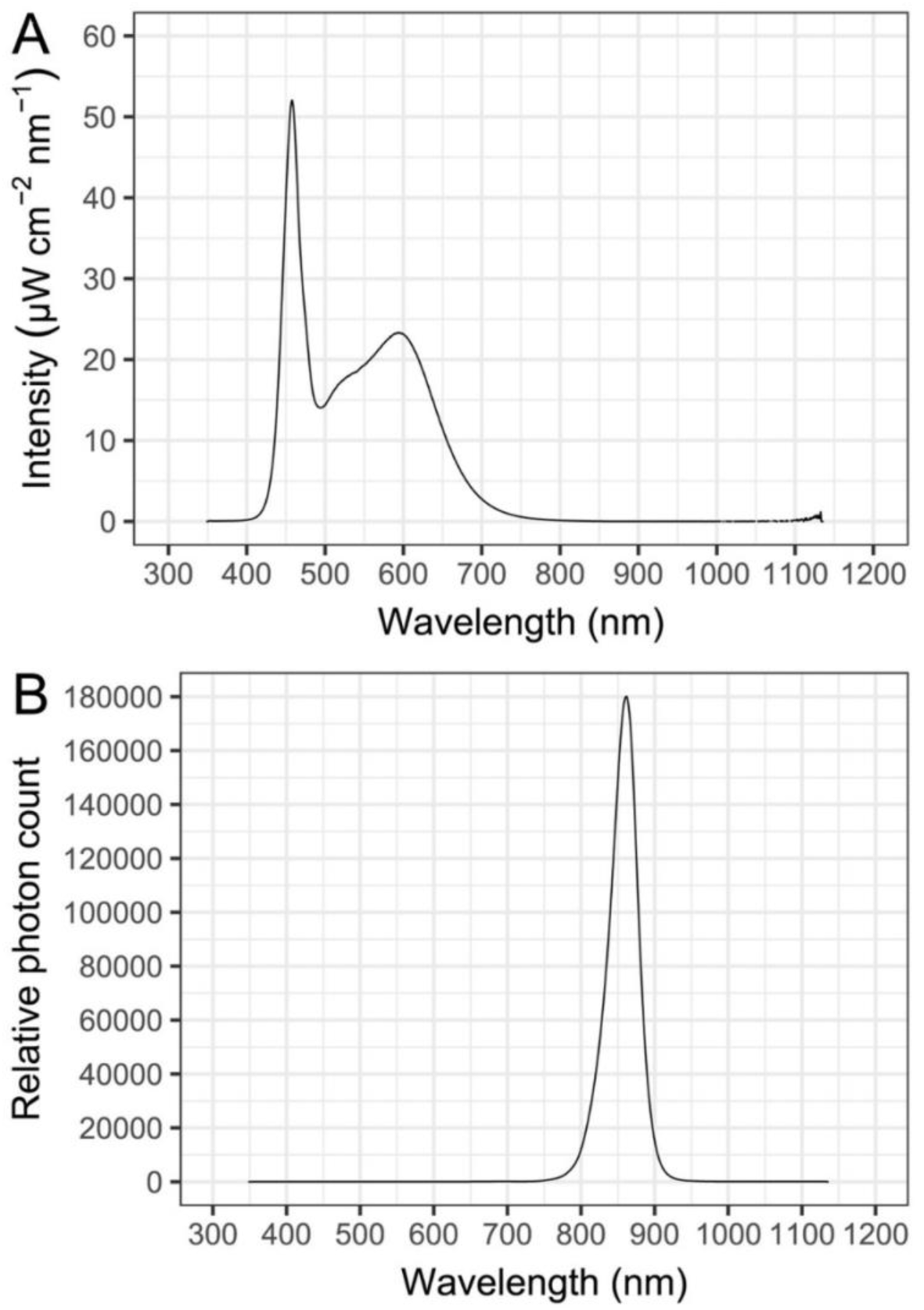
Spectral properties of the light sources used in the experiment. A) Emission spectrum of the LED lamp (FLUVAL Plant Spectrum) used for daytime illumination. Spectra were measured in triplicate, and values are shown as intensity (µW cm⁻² nm⁻¹). The lamp provided 51 W m^−^² at the water surface, positioned ∼30 cm above the sponge explants in the observation chamber. B) Emission spectrum of the infrared (IR) lamp (HD Infrared Waterproof) used for continuous illumination. Spectra were measured in triplicate using an International Light QE Pro spectrometer, and values are shown as relative photon count (normalized to the maximum). The lamp provided 140 W m^−^² at the water surface and was positioned at the same height as the LED light source above the sponge explants in the observation chamber.

**Figure S2.**
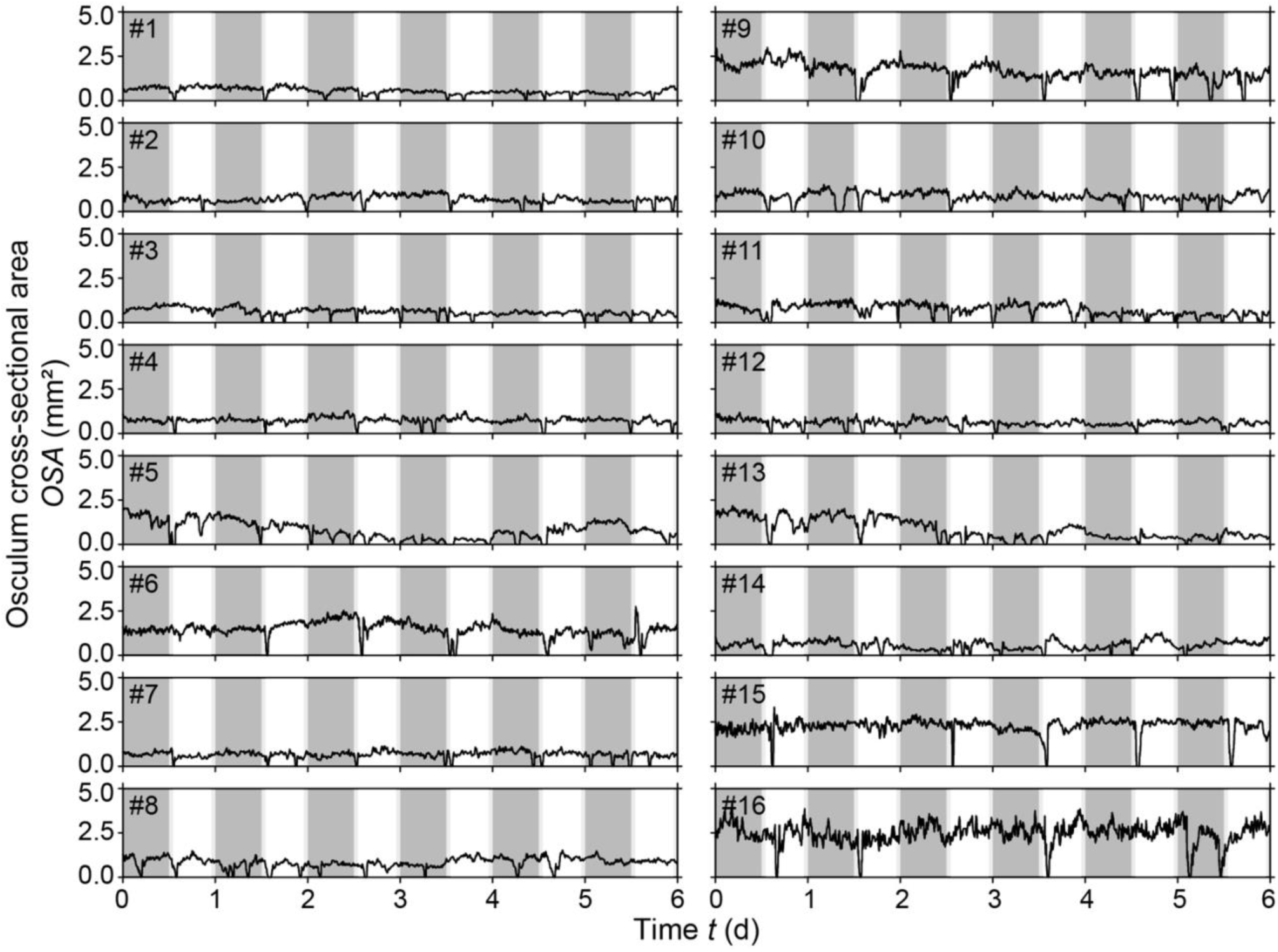
*Halichondria panicea*. Osculum cross-sectional area (*OSA*) of sponge explants (*n* = 16; ID #1‒16) throughout the six-day observation period under controlled light conditions following a 12:12 h dark:light cycle. Dark periods (dark grey areas) and light periods (white areas), including sunrise and sunset (light grey areas), are shown. Contractile behaviour is characterized by complete osculum closure, i.e., reduction of *OSA* to zero, and subsequent expansion of the osculum.

**Figure S3.**
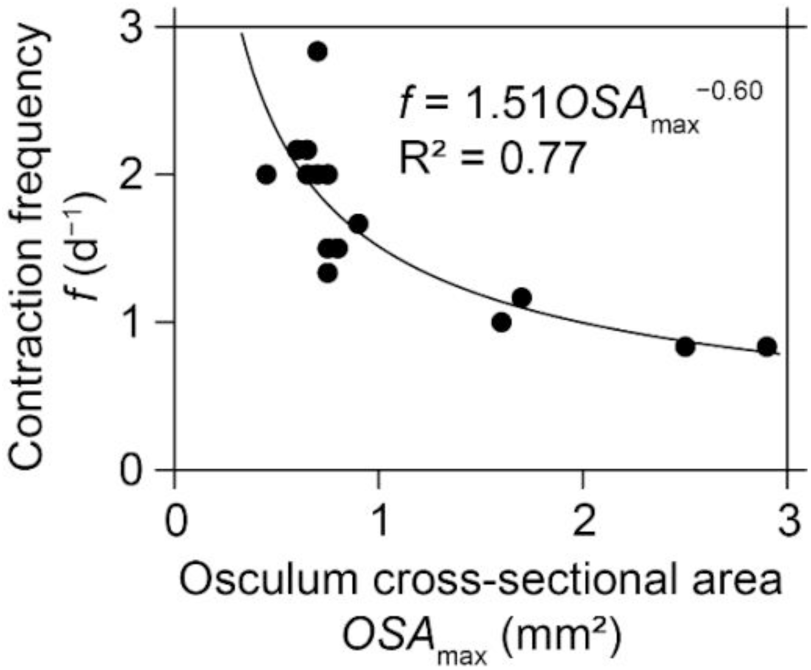
Correlation between osculum size and contraction frequency. Relationship between the contraction frequency and the maximum osculum cross-sectional area of sponge explants (*n* = 16) kept at a 12 h:12 h dark:light cycle. The power function and its equation (LM, t = ‒7.20, p <5×10^‒6^) indicate a 60 % decrease in contraction frequency per 1 mm² increase in osculum cross-sectional area. Data from Table 1.

**Table S1.** Experimental conditions in a seawater flow-through cultivation tank with *Halichondria panicea* explants kept under a 12:12 h dark:light cycle, and hydrography at the sampling site in the inlet to Kerteminde Fjord, Denmark (55°26’59"N, 10°39’40"E). Means ± SD over four replicate experimental sets are shown for the six-day observation period.

| Location | Interval | Time (d) | Temperature (°C) | Salinity (psu) | Chl *a* ($\mu\text{g L}^{-1}$) |
| --- | --- | --- | --- | --- | --- |
| Cultivation tank | Dark | 1 | 15.9±1.9 | 18.9±2.0 | 7.4±1.8 |
| | | 2 | 16.1±1.2 | 19.1±2.2 | 7.5±1.7 |
| | | 3 | 15.7±1.4 | 19.2±2.1 | 7.7±1.6 |
| | | 4 | 16.1±1.1 | 19.8±2.3 | 8.4±1.9 |
| | | 5 | 15.6±1.2 | 20.2±1.9 | 8.6±2.1 |
| | | 6 | 15.4±1.0 | 20.3±1.9 | 8.3±1.3 |
| | Light | 1 | 16.1±1.9 | 19.2±1.9 | 7.3±1.9 |
| | | 2 | 16.1±1.1 | 19.2±2.5 | 7.9±2.2 |
| | | 3 | 15.0±1.5 | 20.1±2.3 | 7.7±1.3 |
| | | 4 | 15.8±1.2 | 20.0±2.1 | 8.5±1.5 |
| | | 5 | 15.6±1.2 | 20.3±2.1 | 8.2±1.6 |
| | | 6 | 15.7±1.1 | 20.2±1.7 | 7.8±1.5 |
| Sampling site | Dark | 1 | 17.3±0.8 | 19.0±1.6 | 8.4±1.4 |
| | | 2 | 17.2±1.4 | 20.1±2.4 | 8.2±1.5 |
| | | 3 | 17.4±1.2 | 21.0±2.8 | 8.4±1.4 |
| | | 4 | 17.5±0.7 | 21.1±2.3 | 8.6±2.0 |
| | | 5 | 17.3±1.1 | 21.3±1.2 | 8.1±1.2 |
| | | 6 | 17.0±0.5 | 21.0±1.9 | 8.5±1.0 |
| | Light | 1 | 17.2±1.2 | 18.8±1.6 | 7.9±1.7 |
| | | 2 | 17.2±0.5 | 20.1±1.6 | 8.0±1.7 |
| | | 3 | 17.3±0.7 | 20.6±1.6 | 7.8±1.7 |
| | | 4 | 17.5±0.5 | 21.0±1.8 | 7.8±1.7 |
| | | 5 | 17.2±0.9 | 20.8±1.6 | 8.0±1.6 |
| | | 6 | 17.1±0.5 | 21.0±1.4 | 8.2±1.3 |

